# Phylogenomics illuminates the backbone of the Myriapoda Tree of Life and reconciles morphological and molecular phylogenies

**DOI:** 10.1101/164616

**Authors:** Rosa Fernández, Gregory D. Edgecombe, Gonzalo Giribet

## Abstract

The interrelationships of the four classes of Myriapoda have been an unresolved question in arthropod phylogenetics and an example of conflict between morphology and molecules. Morphology and development provide compelling support for Diplopoda (millipedes) and Pauropoda being closest relatives, and moderate support for Symphyla being more closely related to the diplopod-pauropod group than any of them are to Chilopoda (centipedes). In contrast, several molecular datasets have contradicted the Diplopoda–Pauropoda grouping (named Dignatha). Here we present the first transcriptomic data including a pauropod and both families of symphylans, allowing myriapod interrelationships to be inferred from phylogenomic data from representatives of all main lineages. Phylogenomic analyses consistently recovered Dignatha with strong support. Taxon removal experiments identified outgroup choice as a critical factor affecting myriapod interrelationships. Diversification of millipedes in the Ordovician and centipedes in the Silurian closely approximates fossil evidence whereas the deeper nodes of the myriapod tree date to various depths in the Cambrian-Early Ordovician, roughly coinciding with recent estimates of terrestrialisation in other arthropod lineages, including hexapods and arachnids. We provide a summary state-of-the-art Myriapoda Tree of Life at the ordinal level, the resolution of which has been improved by phylogenomics.

## Introduction

The evolutionary interrelationships between and within major arthropod groups were subject to much instability in the early years of molecular phylogenetics. Some hypotheses to emerge from that era – such as crustacean paraphyly with respect to Hexapoda (including insects) – have stood the test of time, whereas other have fallen by the wayside. Controversial results were exposed to be artefacts of insufficient amounts of data, flawed analytical methods, or systematic error. In recent years, phylogenomic approaches drawing on vastly expanded gene and taxon coverage, combined with improved analytical approaches, have seen stable, well supported molecular hypotheses being recovered [1-5], and these are increasingly consistent with classical morphological trees.

Transcriptome-based phylogenies drawing on hundreds or thousands of orthologues have assisted with phylogenetic analyses for the major groups of millipedes [6, 7] and centipedes [7, 8] but relationships between the four main myriapod groups have not been as rigorously tested (Fig. 1). A particular limitation is that only Sanger-sequenced data are available for pauropods—a group that lies at the crux of a molecular and morphological conflict within Myriapoda—probably due to their small size, difficulty in finding them, and cryptic behaviour. Analyses based on 62 nuclear protein coding genes [9, 10] underpin a formal taxonomic proposal that Pauropoda is most closely related to Symphyla, a putative clade named Edafopoda [11]. This grouping, also recovered using nuclear ribosomal genes [12] and mitochondrial genomes [13], is highly unexpected from the perspective of morphology because an alternative grouping of Pauropoda and Diplopoda has been widely accepted for more than a century [14-16]. This hypothesis is named Dignatha (=Collifera), referring to the mandibles and first maxillae being the only functional mouthparts, with the limbless postmaxillary segment forming a so-called collum that is not incorporated into the head. Other shared morphological characters include a first maxilla coalesced with a sternal intermaxillary plate, the vas deferens opening to the tips of conical penes between the second trunk leg pair, and the spiracles opening to a tracheal pouch that functions as an apodeme. Early post-embryonic development unites Dignatha based on a motionless pupoid stage immediately after hatching, followed by a hexapodous first free-living stage. The Dignatha hypothesis has also been supported by a few smaller molecular data sets [17], but it has been contradicted by Edafopoda in the analysis of larger data sets [e.g., 11].

**Figure 1.**
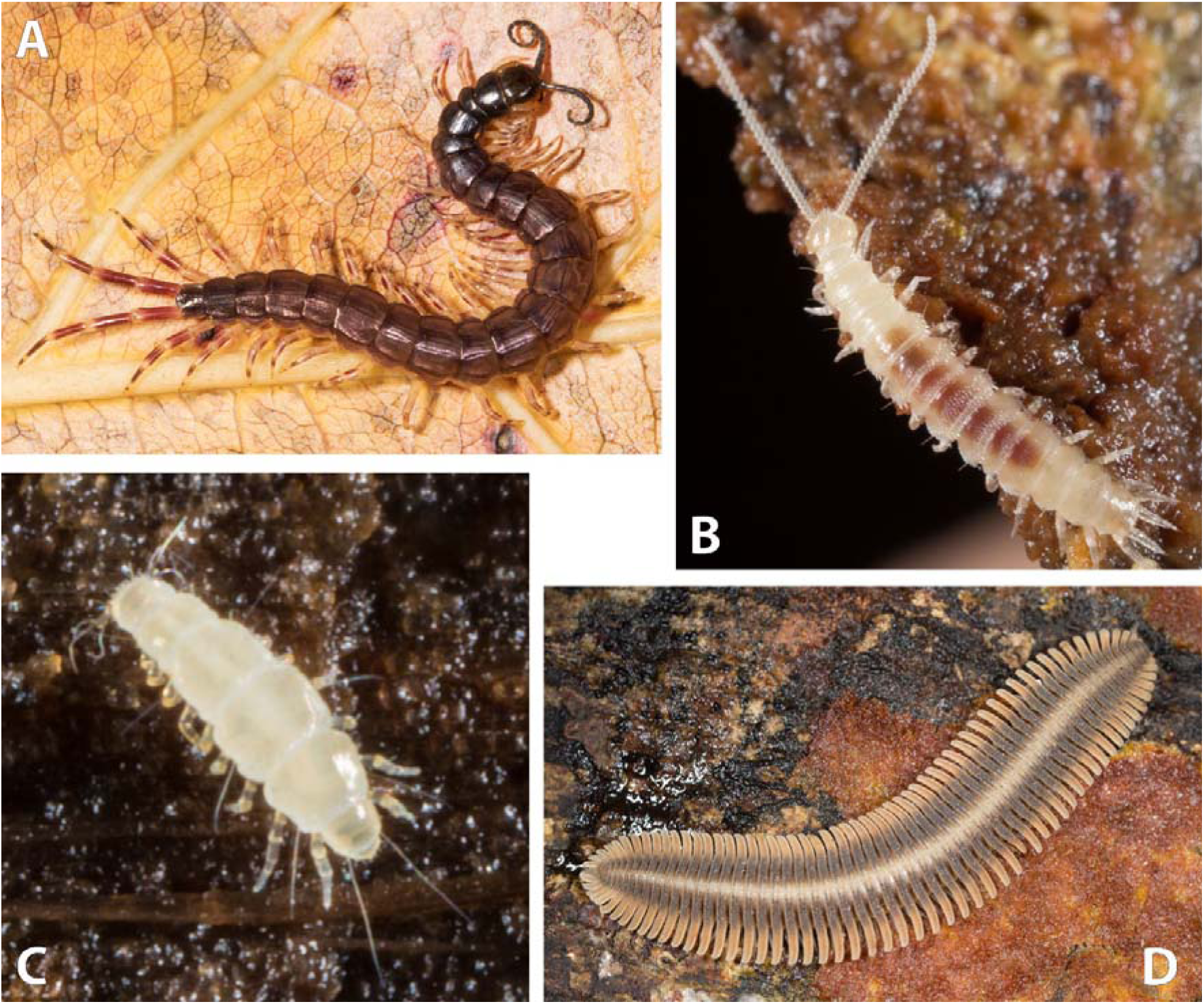
The four main groups of myriapods. A, *Otostigmus* (*Parotostigmus*) *pococki* (Northern Range, Trinidad, Trinidad and Tobago) (Chilopoda, Scolopendromorpha); B, *Hanseniella* sp. (South Island, New Zealand) (Symphyla, Scutigerellidae); C, *Pauropus huxleyi* (Massachusetts, USA) (Pauropoda, Tetramerocerata); and D, *Platydesmus* sp. (La Selva, Costa Rica) (Diplopoda, Platydesmida).

In order to investigate the conflicting support for Dignatha versus Edafopoda – and therefore to shed light on the backbone of the Myriapoda Tree of Life – we present the first transcriptomic data including a pauropod and both families of symphylans. The new data are evaluated in a phylogenomic context specifically designed to test these hypotheses. Furthermore, we expand on previous efforts to date the myriapod phylogenetic tree by coding a morphological character set for the same set of species as sampled transcriptomically as well as key fossil species for their preserved morphological characters in order to estimate the age of diversification within Myriapoda, particularly with reference to the likely timing of terrestrialisation.

## Results & Discussion

### Pauropoda is the sister group to Diplopoda

All the analyses (with the exception of two, in which Pauropoda was attracted to the outgroups, see below) recovered Pauropoda as the sister group to Diplopoda with high support (Fig. 2a,b). Given the unanimity of support for Dignatha/Collifera in morphological studies, this stable, well-supported result reconciles classical morphological studies with molecules. In analyses based on more intensively sampled or closely related outgroups (discussed below), the sister group of Dignatha is Symphyla, together forming the traditional clade Progoneata. The only two analyses not recovering Dignatha (both maximum likelihood analyses not accounting for among-site rate heterogeneity and including the most distant outgroups) positioned Pauropoda at the base of the ingroup due to a long branch attraction artefact (LBA) (Fig. 2c,d). In fact, one of them even recovered Myriapoda as non-monophyletic (Fig. 2d), with the pauropod spuriously clustering within Pancrustacea, highlighting the potential of LBA in this data set. Edafopoda was not recovered in either of these two analyses, as symphylans fall as the sister group of Diplopoda+Chilopoda in both cases (although without strong support in one of the analysis; Fig. 2c). The attraction of symphylans and pauropods as Edafopoda (the only hypothesis exclusively based on molecular information) is therefore probably due to artefacts during phylogenetic reconstruction, as discussed below.

**Figure 2A.**
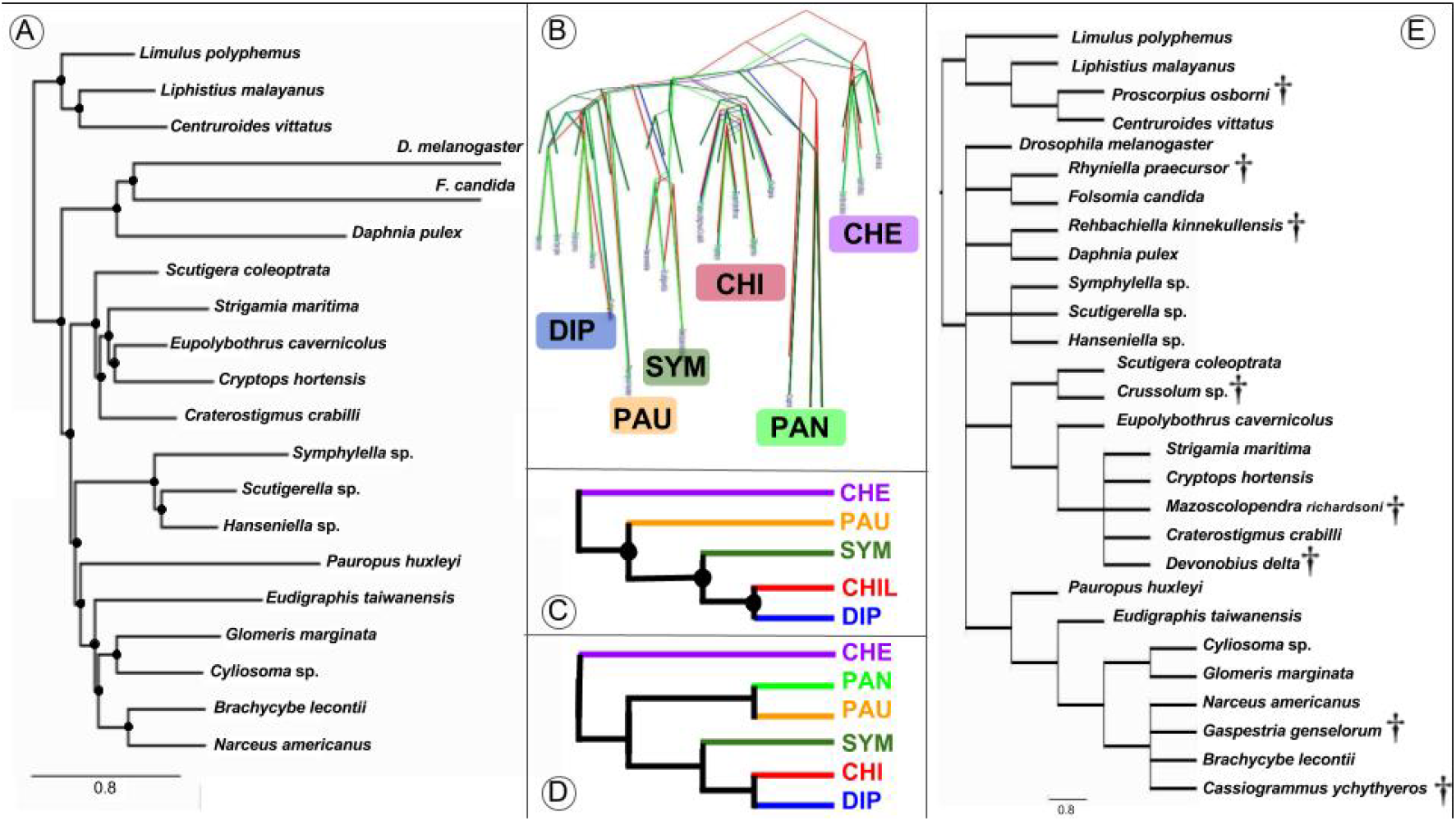
Preferred phylogenetic hypothesis of myriapod interrelationships (PhyloBayes, matrix 1). **2B.** DensiTree visualization of the four most congruent analyseis (PhyloBayes, matrices 2 and 3; PhyML, matrix 3). **2C, 2D.** Main conflicting alternative hypothesis (**2C**, PhyML, matrix 2; **2D**, PhyML, matrix 1). **2E**. Phylogenetic hypothesis of Myriapoda based on 232 morphological characters coded for both extant and extinct species (see Methods for further details); strict consensus of 488 trees of 257 steps; Fossil taxa are identified with a dagger symbol. Black circles in nodes represent high support (> 95% posterior probability, > 90% bootstrap support). CHE: Chelicerata. PAN: Pancrustacea. CHI: Chilopoda. SYM: Symphyla. PAU: Pauropoda. DIP: Diplopoda. Colour codes for each clade are maintained in all figures.

### Outgroup selection impacts on myriapod phylogeny

Despite Dignatha being recovered in most of our analyses, a result not often found in prior molecular analyses, the interrelationships between it and the other two main groups of myriapods varied across analyses, being sensitive to outgroup choice. In the PhyloBayes analyses, matrices 1 and 3 recovered Progoneata, whereas in matrix 2 (in which only the more distantly allied chelicerates were selected as outgroups) symphylans appeared as sister group of the other Myriapoda instead of centipedes, as in the previous cases (Fig. 2a,b). The latter clade formed by these three groups was also recovered in the ML analyses (with or without strong support, though; Fig. 2c,d). This is not the first study in which this result was obtained: in Miyazawa et al. [18] symphylans were likewise recovered as sister group to all other myriapods, followed by pauropods as sister group to millipedes plus centipedes. However, that study was based on just three Sanger sequenced genes, and conflicts with other well resolved nodes in our phylogeny (i.e., Dignatha). The present study and Fernández et al. [7] suggest that outgroup selection is a major factor affecting phylogenetic reconstruction in myriapods. In addition, the latter study found that the most complete matrices were enriched in ribosomal proteins, and both factors strongly compromised the estimated relationships within the ingroup. In the present study, biases from ribosomal proteins were minimized by using a different orthology inference procedure, which ensures that only single copy genes are selected. In spite of this, it remains the case that the selection of only distant outgroups (chelicerates in this case) yields interrelationships of the myriapod classes that are less congruent with morphology than when closer and more comprehensively sampled outgroups are included. This study also highlights the importance of accounting for site-specific heterogeneity (through the CAT-GTR model of PhyloBayes) at least when taxon sampling is not dense for some of the groups, as even when only closer outgroups are included the long-branched pauropod is attracted to the equally long-branched Pancrustacea. The inclusion of more pauropods may alleviate this effect.

### The timing of myriapod diversification

Diversification of Chilopoda (i.e., the basal split in the crown group) is dated to the Early Silurian (Fig. 3), not much earlier than the oldest fossil chilopods in the Late Silurian, these already being representatives of the cholopod crown group. Diversification of Diplopoda dates to the Middle Ordovician (autocorrelated rates)–earliest Silurian (uncorrelated rates). Though this is considerably older than the first millipede body fossils (from the Wenlock Series of the Silurian), it closely approximates the age of trace fossils that have been attributed to Diplopoda and especially compared to locomotion in Penicillata [19, 20]. In contrast, deeper nodes associated with the divergences between myriapod classes are substantially older than available fossil data. No plausible total-group myriapod body fossils are known from the Cambrian, but as in previous studies dating Myriapoda [3, 17], some deep splits are estimated to be of Cambrian age. Diversification of Dignatha is inferred to date to the latest Cambrian-Early Ordovician, Progoneata to the mid-late Cambrian, and Myriapoda to the early-middle Cambrian (auto- and uncorrelated rates, respectively). The shared terrestrial adaptations of all extant myriapods (e.g., tracheae, Malpighian tubules, uniramous trunk limbs) suggest that the common ancestor of each of these estimated Cambrian nodes was terrestrial, coinciding (although being slightly younger) with estimates of terrestrialisation for other arthropod lineages, including arachnids and hexapods [3, 5]. Although the trace fossil record is consistent with amphibious arthropods by the mid Cambrian [21, 22], these molecular estimates for early or middle Cambrian crown-group myriapods continue to pose an unanswered question in arthropod terrestrialisation.

**Figure 3.**
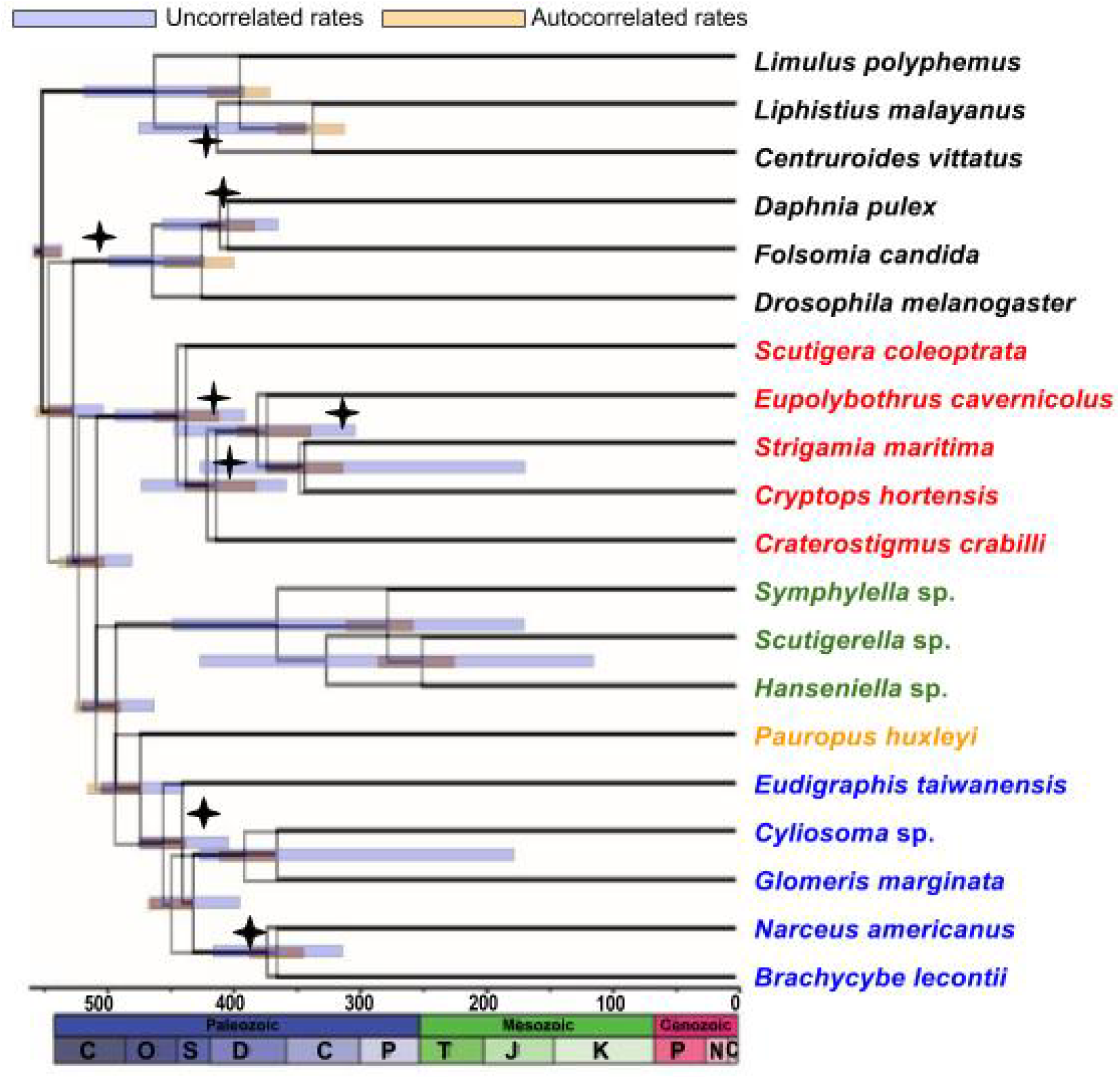
Chronogram of myriapod evolution based on matrix 1 (PhyloBayes analysis) with 95% highest posterior density (HPD) bars for the dating under the uncorrelated (blue) or autocorrelated (pink) model. Nodes that were calibrated with fossils are indicated with a diamond placed at the age of the fossil.

### Towards a fully-resolved Myriapoda Tree of Life

The branching pattern of the four main groups of myriapods has been one of the last unresolved questions in arthropod phylogenetics, together with the interrelationships of the chelicerate orders and the exact sequence of crustaceans that led to the origins of hexapods. With the advent of phylogenomic methods, myriapod phylogeny has attracted attention during the last few years, with several studies devoted to shedding light on the interrelationships of the millipede [6] and centipede [8] orders, and more recently expanding taxon sampling to include most centipede families, most millipede orders and a couple of symphylans [7]. The different analyses of large data matrices combined in all these studies (as well as the current one) have allowed us to discern the main artefacts affecting phylogenetic reconstruction in this group of arthropods and to draw a stable myriapod phylogeny (Fig. 4). The tree, largely congruent with traditional hypotheses based on morphology and development, can now be seen as a well-resolved phylogeny with only a handful of uncertain or untested placements, including the unsampled pauropod order Hexamerocerata, and the unexplored position of the diplopod orders Siphoniulida, Siphonophorida and Siphonocryptida. Further efforts towards including these last missing taxa will set the road towards a complete Myriapoda Tree of Life.

**Figure 4.**
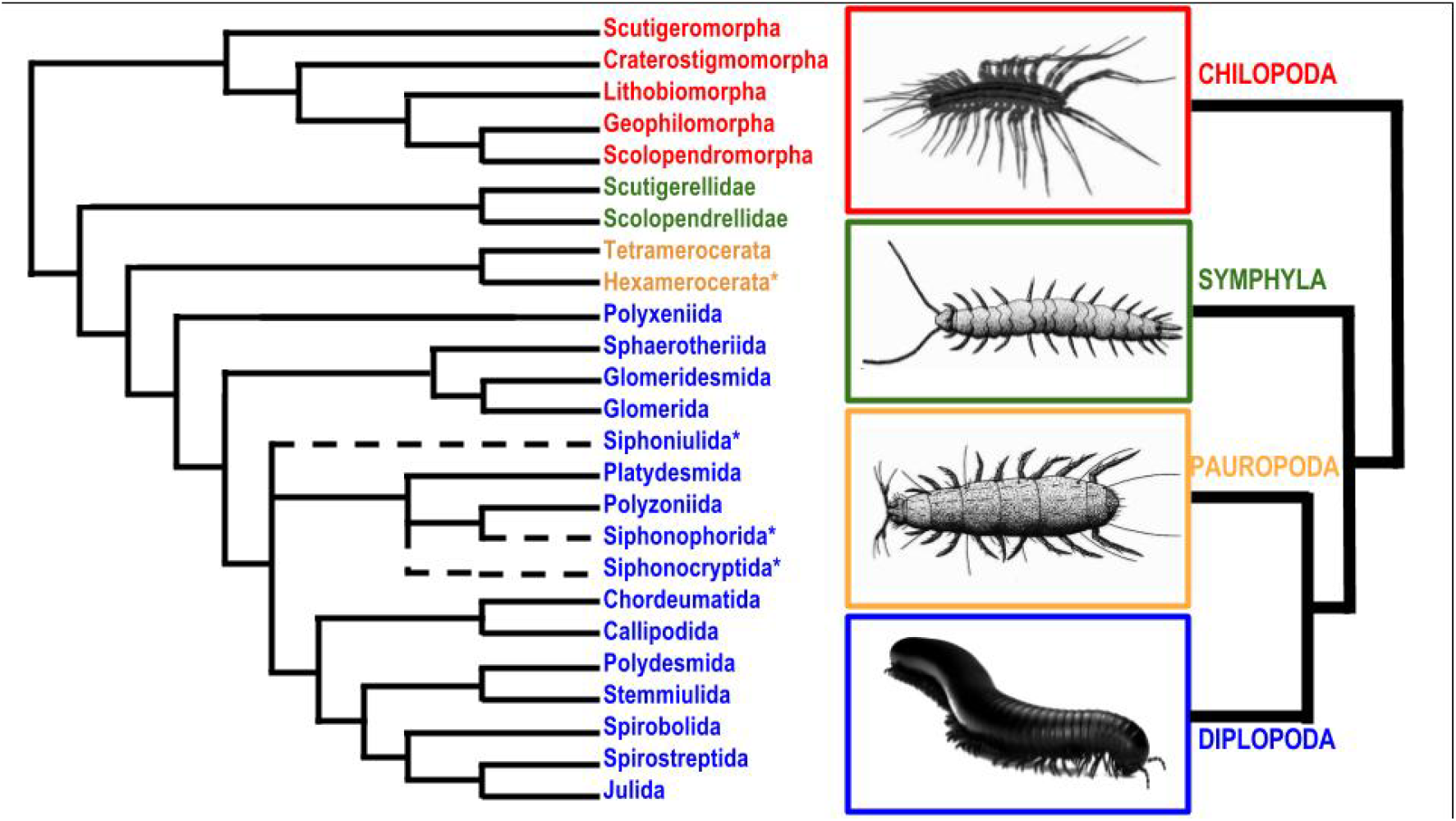
Summary of the Myriapod Tree of Life based as understood by us based on current data. Dashed lines represent unknown placements. Asterisks represent lineages yet to be sequenced with NGS technologies (Hexamerocerata – Pauropoda; Siphoniulida, Siphonophorida, Siphonocryptida – Diplopoda).

## Methods

### Sample collection and molecular techniques

Fourteen species representing the four major groups of myriapods (Chilopoda, Diplopoda, Pauropoda and Symphyla) were included in this study. Building upon previous work [6, 7], our sampling was designed to maximize representation of all groups, including all orders of centipedes, both families of symphylans, the main clades of millipedes, and pauropods. New sequence data were generated from organisms targeted for their instability or lack of representation in prior analyses: a pauropod (*Pauropus huxleyi*) and a symphylan from the family Scolopendrellidae (Scutigerellidae was already represented in earlier studies). Information on sampling localities and accession numbers in the Sequence Read Archive database for each transcriptome can be found in Table 1. The remaining 12 myriapods from Brewer and Bond [6] and our own published data [7] were available from the Sequence Read Archive (SRA). The following taxa were included as outgroups: a crustacean (*Daphnia pulex*), two hexapods (*Drosophila melanogaster, Folsomia candida*), and three chelicerates (*Limulus polyphemus, Liphistius malayanus* and *Centruroides vittatus*). The new sequenced cDNA libraries are accessioned in SRA (Table 1). Tissue preservation and RNA sequencing are as described in Fernández et al. [8]. All data sets included in this study were sequenced with the Illumina platform.

**Table 1.**
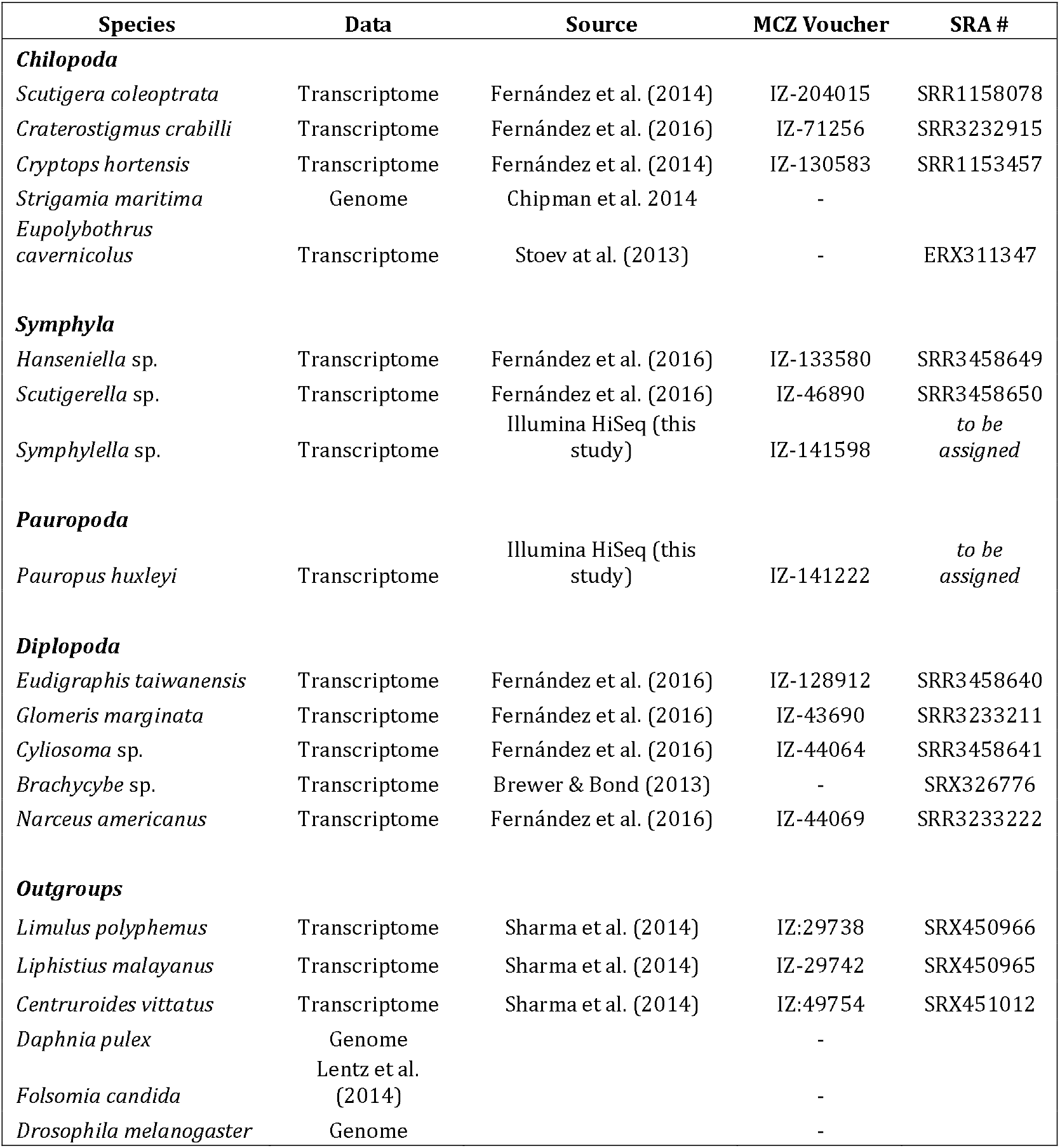
List of taxa included in the present study. Location, Museum of Comparative Zoology (MCZ) and SRA accession numbers are indicated.

### Phylogenomic analyses

Single copy genes in arthropods were identified in our data sets with BUSCO v1 [23] based on hidden Markov model profiles. The homologous genes detected were screened to identify multiple hits (i.e., paralogues). Only one homologue per BUSCO single copy gene was selected in each case, assuming that they were single copy in our samples as well, and therefore orthologs. The genes were parsed from each sample and combined into individual files (i.e., one file per gene) with custom python scripts. Alignment, trimming and concatenation were done as in Fernández et al. [7]. As the selection of outgroups may be critical in resolving myriapod relationships we constructed three matrices with different outgroup composition: matrix 1 (300 genes, 49,576 amino acids), including all outgroups (i.e., chelicerates, hexapods and crustaceans); matrix 2 (229 genes, 49,075 amino acids), only with chelicerate outgroups; and matrix 3 (299 genes, 61,611 amino acids), only with pancrustacean outgroups. In all cases, we selected a high level of gene occupancy to ensure the selection of a relatively large amount of genomic information while minimizing missing data and computational burden (75% gene occupancy in matrix 1 and 88% in matrices 2 and 3). Bayesian analyses were conducted with PhyloBayes MPI 1.4e [24] selecting the site-heterogeneous CAT-GTR model of amino acid substitution [25]. Two independent Markov chain Monte Carlo (MCMC) chains were run for 5000–10,000 cycles. The initial 25% of trees sampled in each MCMC run prior to convergence (judged when maximum bipartition discrepancies across chains were < 0.1) were discarded as burn-in. Convergence of chains was assessed both at the level of the bipartition frequencies (with the command bpcomp) and the summary variables displayed in the trace files (with the command tracecomp). We considered that convergence was achieved when (i) the maximum difference of the frequency of all the bipartitions observed in the chains was < 0.1, and (ii) when the maximum discrepancy observed for each column of the trace file was < 0.1 and the minimum effective size of 100. A 50% majority-rule consensus tree was then computed from the remaining trees. In order to further test for the effect of heterotachy and heterogeneous substitution rates, the matrices where also analysed in PhyML v.3.0.3 implementing the integrated length (IL) approach [26, 27]. In this analysis, the starting tree was set to the optimal parsimony tree and the FreeRate model [28] was selected. Congruence between the different topologies was visualized with DensiTree v2.2.5 [29].

### Molecular Dating

Divergence times for myriapods were estimated through molecular dating, constrained by the position of critical fossils using a morphological data set. Five Palaeozoic and Mesozoic myriapod fossils (two centipedes and three millipedes; Table 2) were included in our morphological data set of 187 characters. We also included three fossil outgroups: the crustacean *Rehbachiella kinnekullensis,* the scorpion *Proscorpius osborni,* and the hexapod *Rhyniella praecursor*. The morphological data set was analysed under parsimony, resulting in 488 trees of 257 steps (Fig. 2e). Absolute dates follow the International Chronostratigraphic Chart v 2015/01. Justifications for age assignments of the fossils (Table 2) follow Wolfe et al. [30]. Divergence dates were estimated using the Bayesian relaxed molecular clock approach as implemented in PhyloBayes v.3.3f [24]. Both an auto-correlated and uncorrelated relaxed clock model were applied to our dataset. The calibration constraints specified above were used with soft bounds [31] under a birth-death prior in PhyloBayes. Two independent MCMC chains were run for 5000–7,000 cycles, sampling posterior rates and dates every 10 cycles. The initial 25% were discarded as burn-in. Posterior estimates of divergence dates were then computed from the remaining samples of each chain.

**Table 2.**
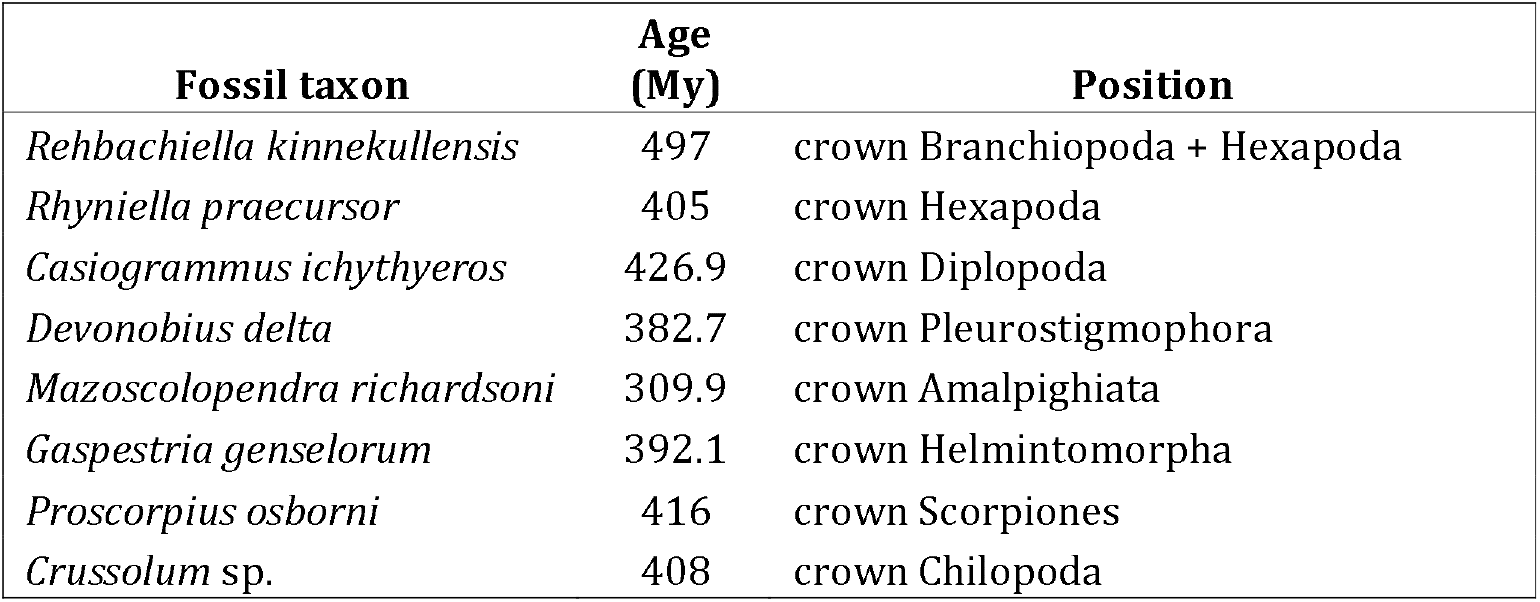
List of fossil taxa selected for the molecular dating analyses. Their age and placement are indicated.

## Author contributions

All authors designed the study. RF and GG collected the samples. RF executed molecular work and data analysis. GDE coded the morphological data. All authors wrote the manuscript.

## Acknowledgements

Miquel Arnedo, Ligia Benavides, Brian Colby and Julia Cosgrove assisted with pauropod collection. This study was mainly supported by internal funds from the Museum of Comparative Zoology, Harvard University and by NSF grant DEB-1457539 to GG, which funded RF.

## Data availability statement

All data sets have been deposited to the NCBI SRA data base. Accession numbers are listed in Table 1.

